# The ATF6β-calreticulin axis promotes neuronal survival under endoplasmic reticulum stress and excitotoxicity

**DOI:** 10.1101/2021.02.01.429116

**Authors:** Dinh Thi Nguyen, Thuong Manh Le, Tsuyoshi Hattori, Mika Takarada-Iemata, Hiroshi Ishii, Jureepon Roboon, Takashi Tamatani, Takayuki Kannon, Kazuyoshi Hosomichi, Atsushi Tajima, Shusuke Taniuchi, Masato Miyake, Seiichi Oyadomari, Shunsuke Saito, Kazutoshi Mori, Osamu Hori

**Affiliations:** Department of Neuroanatomy, Graduate School of Medical Sciences, Kanazawa University, Kanazawa, Japan; Department of Bioinformatics and Genomics, Graduate School of Advanced Preventive Medical Sciences, Kanazawa University, Kanazawa, Japan; Division of Molecular Biology, Institute for Genome Research, Institute of Advanced Medical Sciences, Tokushima University, Tokushima, Japan; Department of Biophysics, Graduate School of Science, Kyoto University, Kyoto, Japan

**Author notes:** Corresponding author: Dr. Osamu Hori, Department of Neuroanatomy, Kanazawa University Graduate School of Medical Sciences, 13-1 Takara-Machi, Kanazawa City, Ishikawa 920-8640, Japan.

**Keywords:** neurodegeneration, Ca^2+^ homeostasis, ER stress

## Abstract

While ATF6α plays a central role in the endoplasmic reticulum (ER) stress response, the function of ATF6β is largely unknown. Here, we demonstrate that ATF6β is highly expressed in the hippocampus of the brain, and specifically regulates the expression of calreticulin, a molecular chaperone in the ER with a high Ca^2+^-binding capacity. Calreticulin expression was reduced to ~50% in the central nervous system of *Atf6b^−/−^* mice, and restored by ATF6β. Analysis using cultured hippocampal neurons revealed that ATF6β deficiency reduced Ca^2+^ stores in the ER and enhanced ER stress-induced death, which was rescued by ATF6β, calreticulin, Ca^2+^-modulating reagents such as BAPTA-AM and 2-APB, and ER stress inhibitor salubrinal. *In vivo*, kainate-induced neuronal death was enhanced in hippocampi of *Atf6b^−/−^* and *Calr^+/−^* mice, and restored by 2-APB and salubrinal. These results suggest that the ATF6β-calreticulin axis plays a critical role in the neuronal survival by improving Ca^2+^ homeostasis under ER stress.

## Introduction

The endoplasmic reticulum (ER) is an intracellular organelle in which secretory proteins and lipids are synthesized, and intracellular Ca^2+^ is stored. However, recent studies demonstrated that a stress response occurs in the ER. When cells are exposed to specific conditions such as impaired Ca^2+^ homeostasis, energy shortage, and increased protein synthesis, unfolded proteins accumulate in the ER, leading to a condition generally termed ER stress (Bukau *et al.*, 2006). In the central nervous system (CNS), pathological situations such as brain ischemia, neurodegeneration, excitotoxicity and demyelination are tightly associated with ER stress (Sokka *et al.*, 2007; Sprenkle *et al.*, 2017; Thiebaut *et al.*, 2019). Cells can respond to ER stress by activating the unfolded protein response (UPR). There are at least three transducers of the UPR, namely, protein kinase R (PKR)-like ER kinase (PERK), inositol-requiring enzyme 1 (IRE1), and activating transcription factor 6 (ATF6) (Mori, 2009; Walter & Ron, 2011). Among these, ATF6 is responsible for induction of the major molecular chaperones in the ER, such as glucose-regulated protein 78 (GRP78) and glucose-regulated protein 94 (GRP94), in addition to several ER-associated degradation components such as homocysteine-responsive ER-resident ubiquitin-like domain member 1 protein and ER degradation enhancing α mannosidase (Yamamoto *et al.*, 2007; Mori, 2009). In mammals, there are two subtypes of ATF6, called ATF6α and ATF6β. Both molecules are type II transmembrane proteins in the ER and translocate to the Golgi apparatus for cleavage upon ER stress. Although ATF6α plays a dominant role in the transcriptional activation in response to ER stress, ATF6α/β-mediated adjustment of chaperone levels to meet the increased demands in the ER is essential for the development of the notochord (Yamamoto *et al.*, 2007; Ishikawa *et al.*, 2013).

We previously reported that ATF6α contributes to both neuronal survival and glial activation in different neuropathological situations. Deletion of *Atf6α* or that of a downstream molecular chaperone, *Hyou1*, sensitizes hippocampal neurons to glutamate-induced toxicity most likely via Ca^2+^ overload and neuronal hyperactivity *in vivo* (Kitao *et al.*, 2001; Kezuka *et al.*, 2016). *Atf6α* deficiency is also associated with reduced astroglial activation and glial scar formation in mouse models of Parkinson’s disease (Hashida *et al.*, 2012) and stroke (Yoshikawa *et al.*, 2015), respectively, both of which are associated with an enhanced level of neuronal death. By contrast, in experimental autoimmune encephalomyelitis mice, a model of multiple sclerosis, and cultured microglia, *Atf6α* deficiency suppresses microglial activation, clinical symptoms and demyelination via a mechanism involving rapid degradation of NF-κB p65 by the proteasome (Ta *et al.*, 2016).

In contrast with the role of ATF6α in the ER stress response/UPR and ER stress-related pathophysiologies, the function of ATF6β is largely unknown, especially in the CNS. Transcriptional activity of ATF6β is reportedly much weaker than that of ATF6α (Haze *et al.*, 2001). However, recent reports demonstrated that ATF6α and ATF6β may have overlapping and differential functions in the mouse heart (Lynch *et al.*, 2012; Correll *et al.*, 2019). We therefore sought to investigate the expression and possible roles of ATF6β in the CNS under normal and ER stress conditions. Here, we demonstrate that calreticulin (CRT), a molecular chaperone in the ER with a high Ca^2+^-binding capacity, is a unique target of ATF6β in the CNS, and the ATF6β-CRT axis plays a critical role in the neuronal survival by improving Ca^2+^ homeostasis under ER stress.

## Results

### Expression of ATF6β in the CNS and other tissues

We first verified the tissue distribution of ATF6β in mice. Quantitative real-time PCR (qRT-PCR) revealed that *Atf6b* mRNA was broadly expressed, but was most highly expressed in the hippocampus of the brain among the tissues analyzed (Figure 1A). Further analysis of cultured cells revealed that expression of *Atf6b* mRNA was higher in hippocampal neurons than in cortical neurons and astrocytes under normal conditions (Figure 1B). Consistently, *in situ* hybridization revealed that *Atf6b* mRNA was highly expressed in hippocampal neurons (Figure 1C). These patterns were in contrast with those of *Atf6a* mRNA, which was more ubiquitously expressed (Figure S1 A, B). There was no significant difference in *Atf6b* mRNA levels between male and female mice (Figure S1 C).

**Figure 1.**
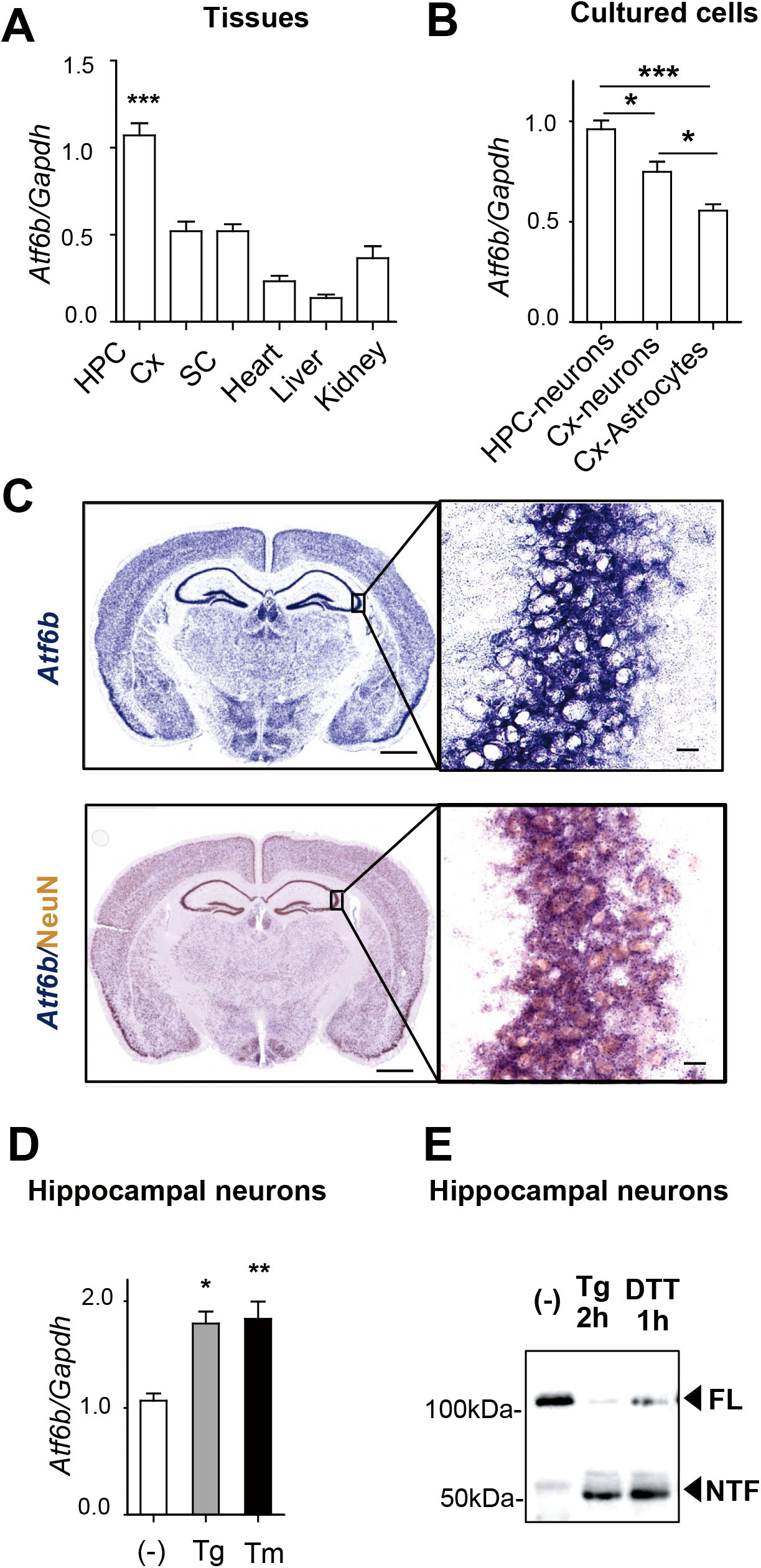
Expression and activity of ATF6β. A, B, Expression of *Atf6b* mRNA in normal tissues (n=5 mice) (A) and in cultured cells (n=3–8) (B). HPC: hippocampus, Cx: cerebral cortex, SC: spinal cord. Total RNA was isolated from the indicated samples and qRT-PCR was performed. Data are shown as mean ± SEM. *p < 0.05, ***p < 0.001 by a one-way ANOVA followed by the Tukey test. C, *In situ* hybridization (upper panel) and *in situ* hybridization-immunohistochemistry (lower panel) of *Atf6b* mRNA in the normal brain. Images in the right panels are enlarged views of the CA3 area. Scale bars: 200 μm (left panels) and 25 μm (right panels). Typical images from three independent experiments are shown. D, qRT-PCR analysis of expression of *Atf6b* mRNA in primary hippocampal neurons under ER stress. Cells were treated with Tg (300nM) or Tm (1μg/ml) for 8 h and then qRT-PCR was performed. n=3. Data are shown as mean ± SEM. *p < 0.05, **p < 0.01 by a one-way ANOVA followed by the Tukey test. E, Activation of ATF6β by ER stress in primary hippocampal neurons. Cells were treated with Tg (300nM) for 2h or DTT for 1h. Extracted proteins were subjected to western blotting. The typical data from two independent experiments are shown. FL: full length, NTF: N-terminal fragment

We next analyzed expression of *Atf6b* mRNA under ER stress. Treatment of cultured hippocampal neurons with the ER stressors tunicamycin (Tm) and thapsigargin (Tg) significantly increased expression of *Atf6b* mRNA (1.5–2-fold increase) (Figure 1D), although these increases were smaller than those in expression of *Atf6a* mRNA (5–6.5-fold increase) (Figure S1 D). At the protein level, both the full-length 110 kDa protein (FL) and a cleaved N-terminal 60kDa fragment (NTF) of ATF6β were detected in primary hippocampal neurons. The level of this fragment was low under normal conditions, but increased as early as 2h after Tg treatment or 1h after dithiothreitol (DTT) treatment, the latter also causes ER stress. These results suggest that ATF6β functions in neurons especially under ER stress.

### Calr is a unique target gene of ATF6β in the CNS

To identify downstream molecules of ATF6β in the CNS, RNA-sequencing was performed using hippocampal brain samples from wild-type (WT) and *Atf6b^−/−^* mice. A total of 55,531 genes were examined. We filtered genes in two ways. When filtering genes with FPKM values in WT mice higher than 10 and q values smaller than 0.05, only 2 downregulated genes and 4 upregulated genes were identified in *Atf6b^−/−^* mice (Table 1). Although expression of *Atf6b* mRNA was observed to some extent in *Atf6b^−/−^* mice, this may be due to the presence of the 5’ *Atf6b* transcript with exon 1-9 in these mice, in which exon 10 and 11 were deleted by homologous recombination (Yamamoto *et al.*, 2007) (Figure S2). Besides *Atf6b*, only *Calr*, which encodes CRT, a molecular chaperone in the ER with a high Ca^2+^-binding capacity, was downregulated in *Atf6b^−/−^* mice (Table 1). By contrast, in case filtering genes with FPKM values in WT mice higher than 10 and p values smaller than 0.05, 22 downregulated genes and 27 upregulated genes were identified in *Atf6b^−/−^* mice (Table S1). *Calr* was again identified as a gene downregulated in *Atf6b^−/−^* mice, and interestingly, six ER stress-responsive genes, namely, *Hpsa5* (GRP78), *Pdia4* (ERP72), *Dnajb11*, *Atf4*, *Wfs1,* and *P4ha1* were upregulated in *Atf6b^−/−^* mice (Table S1 right columns). RNA-sequencing also indicated that expression of *Calr* was highest among the major molecular chaperones in the ER in WT brains (Table S2). Taken together with previous reports demonstrating possible roles of ATF6β in the expression of molecular chaperones in the ER (Yamamoto *et al.*, 2007; Ishikawa *et al.*, 2013; Correll *et al.*, 2019), we decided to focus on ATF6β-CRT axis in further experiments.

**Table 1.** Differentially expressed genes in *Atf6b^−/−^* brain (q<0.05)

| Downregulated genes in *Atf6b*<sup>-/-</sup> brain |  |  |  |  | Upregulated genes in *Atf6b*<sup>-/-</sup> brain |  |  |  |  |
| --- | --- | --- | --- | --- | --- | --- | --- | --- | --- |
| Gene | WT | *Atf6b*<sup>-/-</sup> | log2 (fold-change) | q-value | Gene | WT | *Atf6b*<sup>-/-</sup> | log2 (fold-change) | q-value |
| *Atf6b* | 31.633 | 7.66511 | -2.04505 | 0.0465921 | *Gm47283* | 13.354 | 43.6105 | 1.70741 | 0.0465921 |
| *Calr* | 203.013 | 123.236 | -0.720147 | 0.0465921 | *Igf2* | 19.0054 | 39.9251 | 1.07089 | 0.0465921 |
|  |  |  |  |  | *Ptgds* | 452.8 | 942.637 | 1.05783 | 0.0465921 |
|  |  |  |  |  | *Mgp* | 20.9161 | 35.9567 | 0.781646 | 0.0465921 |

Consistent with the RNA-sequencing data, RT-qPCR using hippocampi of WT, *Atf6a ^−/−^*, and *Atf6b^−/−^* mice revealed that expression of *Calr* mRNA was reduced to ~50% in *Atf6b^−/−^* brains, but not in *Atf6a^−/−^* brains (Figure 2A). This was in contrast with expression of *Hspa5* (GRP78) mRNA, which was reduced in *Atf6a^−/−^* brains, but increased in *Atf6b^−/−^* brains (Figure 2A). Similar differences in expression of *Hsp90b1* (GRP94) mRNA were observed, although these were not significant. Expression of *Canx*, which encodes calnexin, another molecular chaperone in the ER with similarities to CRT, was unaffected by *Atf6a* and *Atf6b* deficiency (Figure 2A).

**Figure 2.**
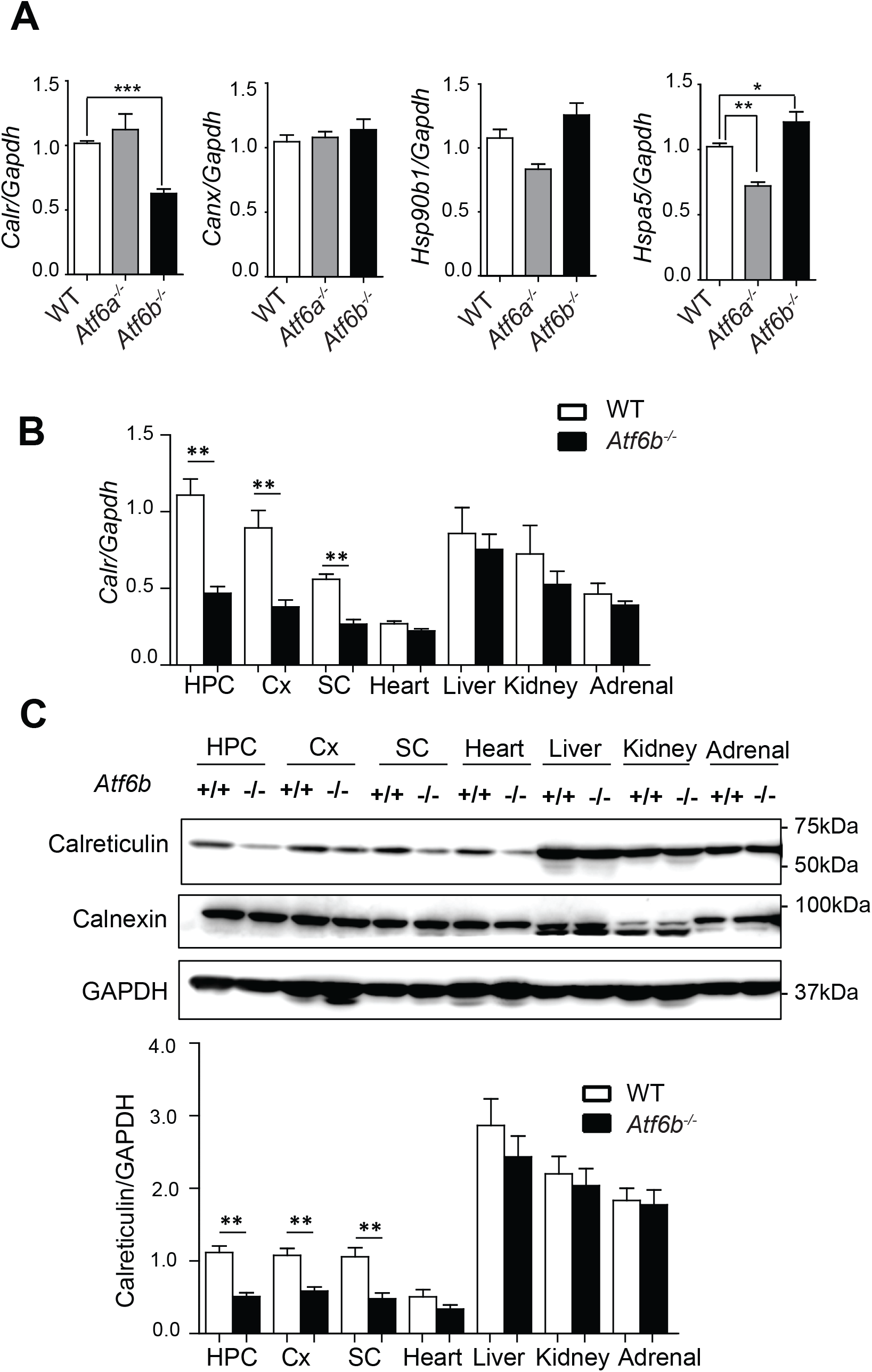
Reduces CRT expression in the CNS of *Atf6b^−/−^* mice. A, Expression of molecular chaperones in the ER in WT, *Atf6a ^−/−^* and *Atf6b^−/−^* hippocampi. Total RNA was isolated from the CA3 region of the hippocampus and qRT-PCR was performed. n=5–6 mice. Data are shown as mean ± SEM. *p < 0.05, **p < 0.01, ***p < 0.001 by a one-way ANOVA followed by the Tukey test. B, Expression of *Calr* mRNA in WT and *Atf6b^−/−^* tissues. Total RNA was isolated from the indicated tissues of each mouse and qRT-PCR was performed. n=5 mice. Data are shown as mean ± SEM. **p < 0.01 by the Mann-Whitney U test. HPC: hippocampus, Cx: cerebral cortex, SC: spinal cord. C, Expression of CRT protein in WT and *Atf6b^−/−^* tissues. Protein samples were extracted from the indicated tissues of WT and *Atf6b^−/−^* mice, and subjected to western blotting. n=5–7 mice. Data are shown as mean ± SEM. **p < 0.01 by the Mann-Whitney U test.

The effect of *Atf6b* deletion on CRT expression was next analyzed in different tissues under normal conditions (Figure 2B, C). Both qRT-PCR (Figure 2B) and western blotting (Figure 2C) revealed that CRT expression was significantly lower in the CNS, but not in other tissues tested, in *Atf6b^−/−^* mice than in WT mice.

### The role of ATF6β in CRT promoter activity

To analyze the role of ATF6β in CRT expression at the promoter level, reporter assays were performed with a chloramphenicol acetyltransferase (CAT) plasmid containing 1763bp and 415bp of the mouse CRT promoters, pCC1 and pCC3, respectively (Waser *et al.*, 1997) and luciferase plasmids containing 459bp of the WT (huCRT(wt)) or mutant (huCRT(mut)) human CRT promoter, with the latter containing mutated sequences of two ER stress-responsive elements (ERSEs) (Yoshida *et al.*, 1998) (Figure 3A). Upon transfection of pCC1 or pCC3, promoter activity was lower in *Atf6b^−/−^* hippocampal neurons (59% in pCC1 and 58% in pCC3, respectively) than in WT hippocampal neurons (Figure 3B). Overexpression of ATF6β cDNA restored the promoter activity in *Atf6b^−/−^* neurons (103% in pCC1 and 132% in pCC3). Similarly, upon transfection of huCRT(wt), CRT promoter activity was lower (59%) in *Atf6b^−/−^* neurons than in WT neurons, and overexpression of ATF6β cDNA restored promoter activity to 87% in *Atf6b^−/−^* neurons (Figure 3C). Interestingly, overexpression of ATF6α cDNA restored CRT promoter activity to a greater extent in the same condition (187% in pCC1, 201% in pCC3, and 109% in huCRT(wt)) in *Atf6b^−/−^* neurons. By contrast, upon transfection of huCRT(mut), CRT promoter activity was not detected in either WT or *Atf6b^−/−^* neurons and was not restored at all by overexpression of ATF6β or ATF6α cDNA (Figure 3C), suggesting that ERSEs are essential for ATF6β-mediated CRT promoter activation.

**Figure 3.**
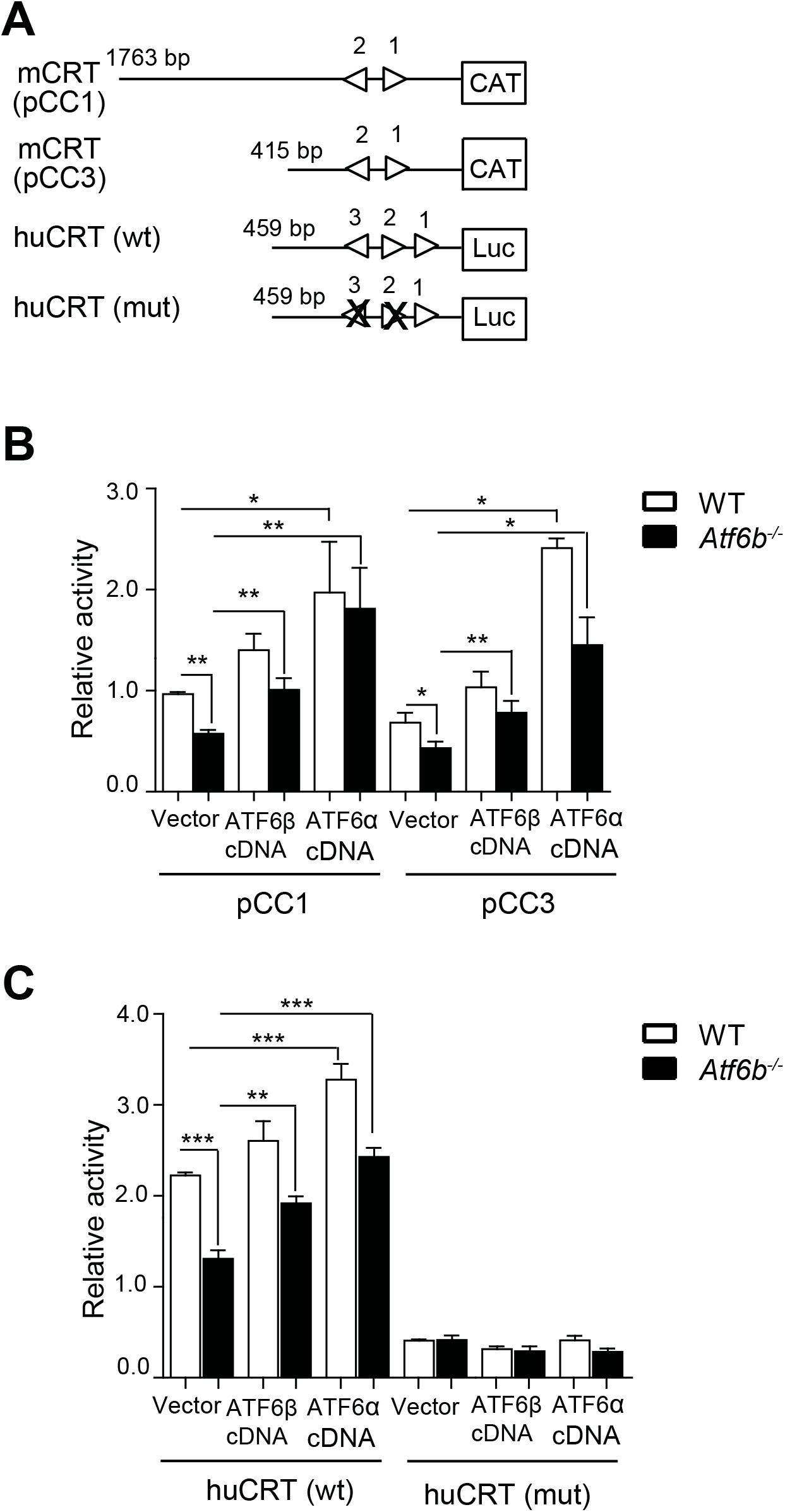
Reduced CRT promoter activity in Atf6b-deleted neurons. A, Schematic representation of the promoters used. Triangles indicate the locations and orientations of ERSE motifs that completely or considerably match the consensus CCAATN9CCACG (Yoshida *et al.*, 1998). Numbers indicate nucleotide positions from transcription start site. ERSE2 and ERSE3 of the human CRT promoter were disrupted by mutating their sequences (marked by crosses). B, C, Reporter assays using cultured hippocampal neurons. The CAT ELISA and luciferase assay were performed using cells transfected with the mouse CRT promoter pCC1 and pCC3 (B), or with the human CRT promoter huCRT (C). n=4. Data are shown as mean ± SEM. *p < 0.05, **p < 0.01, ***p < 0.001. by a two-way ANOVA followed by the Bonferroni test. Note that both ATF6α and ATF6β cDNAs increased CRT promoter activity in *Atf6b^−/−^* cells.

Similar results were obtained in mouse embryonic fibroblasts (MEFs) transfected with pCC1, huCRT(wt), or huCRT(mut), although ATF6β cDNA seemed to have slightly higher restoring effects than ATF6α cDNA in *Atf6b^−/−^* cells (Figure S3). In contrast to CRT promoter activity, GRP78 promoter activity, which was measured using a luciferase plasmid containing 132 bp of the human GRP78 promoter (Yoshida *et al.*, 1998), did not differ between WT and *Atf6b^−/−^* neurons under normal or ER stress conditions (data not shown). These results suggest that ATF6β specifically enhances CRT promoter activity in a similar manner to ATF6α via ERSE motifs.

### Effect of ATF6β deletion on expression of molecular chaperones in the ER in primary hippocampal neurons

The effect of ATF6β deletion on expression of molecular chaperones in the ER was next examined under normal and ER stress conditions using WT and *Atf6b^−/−^* hippocampal neurons. Consistent with the results obtained using mouse tissues under normal conditions, RT-qPCR revealed that *Calr* expression was significantly lower in *Atf6b^−/−^* neurons than in WT neurons under both control and ER stress conditions, with the latter induced by Tg (Figure 4A upper row) and Tm (Figure 4A lower row). By contrast, expression of other molecular chaperones in the ER such as *Canx* (calnexin), *Hsp90b1* (GRP94), and *Hspa5* (GRP78) was temporally lower in *Atf6b^−/−^* neurons than in WT neurons after stimulation with Tg (Figure 4A upper row) or Tm (Figure 4A lower row). Similarly, western blot analysis revealed that expression of CRT protein was constitutively lower in *Atf6b^−/−^* neurons than in WT neurons, while protein expression of other molecular chaperones in the ER was similar in *Atf6b^−/−^* and WT neurons under both normal and ER stress conditions (Figure 4B).

**Figure 4.**
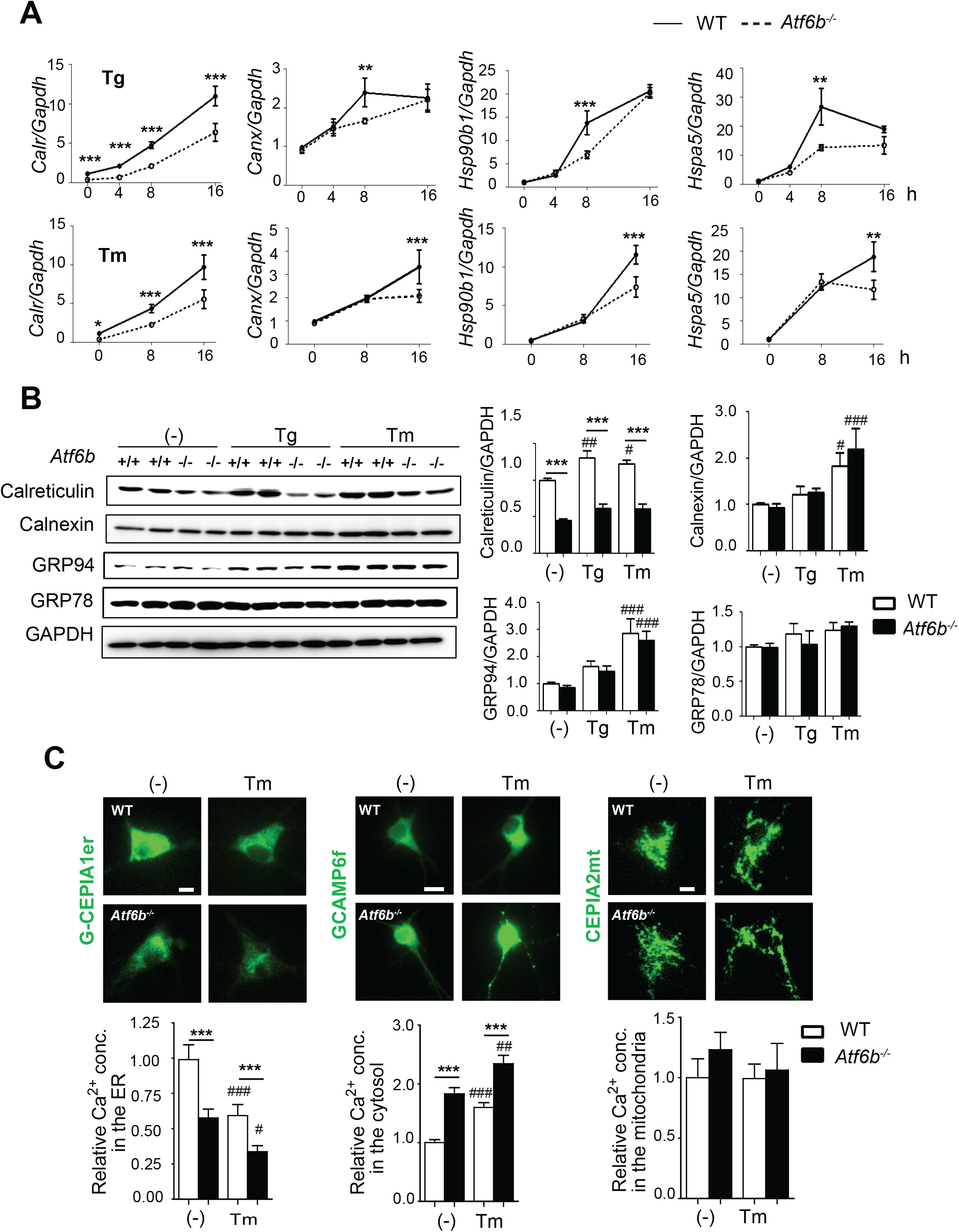
Expression of molecular chaperones in the ER and Ca^2+^ homeostasis in *Atf6b^−/−^* hippocampal neurons. A, Total RNA was isolated from WT and *Atf6b^−/−^* hippocampal neurons (n=3–8), and qRT-PCR was performed with the indicated primers. Data are shown as mean ±SEM. *p < 0.05, **p < 0.01, ***p<0.001 between two genotypes by a two-way ANOVA followed by the Bonferroni test. Note that expression of *Calr* mRNA was significantly lower in *Atf6b^−/−^* neurons than in WT neurons under both normal and ER stress conditions. B, Protein samples extracted from WT and *Atf6b^−/−^* hippocampal neurons exposed to control or ER stress conditions for 16h (n=5–6) were analyzed by western blotting using antibodies against the indicated proteins. Data are shown as mean ± SEM. ***p < 0.001 between two genotypes and # p < 0.05, ## p < 0.01, ### p<0.001 compared to normal conditions by a two-way ANOVA followed by the Bonferroni tests. Note that expression of CRT protein was significantly lower in *Atf6b^−/−^* neurons than in WT neurons under both normal and ER stress conditions. C, Ca^2+^ measurement. Ca^2+^ levels in the ER, cytosol and mitochondria of hippocampal neurons were measured using G-CEPIA1er, GCaMP6f and CEPIA2mt, respectively under normal and ER stress (Tm for 3h) conditions. n=60–250 cells in each condition from two independent experiments. Data are shown as mean ± SEM. ***p < 0.001 between two genotypes and # p < 0.05, ## p < 0.01, ### p<0.001 compared to normal conditions by a two-way ANOVA followed by the Bonferroni tests.

Finally, gene-silencing experiments were performed to exclude the possibility that the effect of *Atf6b* deletion on CRT expression was indirect due to the long-term absence of *Atf6b*. Transfection of Neuro 2a cells with two sets of ATF6β-targeting siRNAs (ATF6β-siRNA1 and ATF6β-siRNA2) reduced *Atf6b* expression to 30% and 43% and reduced *Calr* expression to 62% and 66%, but did not affect *Hspa5* (GRP78) expression, compared with that in control-siRNA-transfected cells (Figure S5 A).

### Effect of ATF6β deletion on Ca^2+^ homeostasis in primary hippocampal neurons

Consistent with the reduced level of *Calr* expression in *Atf6b^−/−^* neurons, Ca^2+^ levels in the ER, which were measured by the green fluorescence-Ca^2+^-measuring organelle-entrapped protein indicator 1 in the ER (G-CEPIA1er) (Suzuki *et al.*, 2014), were lower in the in *Atf6b^−/−^* neurons under both normal and ER stress conditions (Figure 4C left). By contrast, Ca^2+^ levels in the cytosol, which were measured by GFP-based Ca^2+^ calmodulin probe 6f (GCaMP6f)(Chen *et al.*, 2013) were higher in *Atf6b^−/−^* neurons under both normal and ER stress conditions (Figure 4C middle). Ca^2+^ levels in the mitochondria, which were measured by CEPIA2 in the mitochondria (CEPIA2mt)(Suzuki *et al.*, 2014), were at similar levels between two genotypes under both normal and ER stress conditions (Figure 4C right).

### The neuroprotective role of the ATF6β-CRT axis against the ER stress-induced neuronal death

To evaluate whether ATF6β has a neuroprotective role against ER stress, WT and *Atf6b^−/−^* hippocampal neurons were treated with Tg or Tm and the cell death/survival was evaluated in two ways. Staining of living and dead cells with the fluorescent dyes, calcein-AM (green) and ethidium homodimer-1 (EthD-1, red), respectively, revealed that almost all cells are alive in normal conditions. Treatment with ER stressors induced death and reduced viability of both WT and *Atf6b^−/−^* neurons, but this effect was more pronounced in *Atf6b^−/−^* neurons (Figure 5A, Figure S4). Consistently, immunocytochemistry using an antibody against the apoptosis marker cleaved caspase-3 (red) and neuronal marker βIII tubulin (green) indicated the caspase-3 activation and loss of βIII tubulin expression were higher in *Atf6b^−/−^* neurons than in WT neurons under ER stress (Figure 5B). Similarly, transient silencing of *Atf6b* gene using siRNA enhanced ER stress-induced death of Neuro 2a cells (Figure S5 B, C).

**Figure 5.**
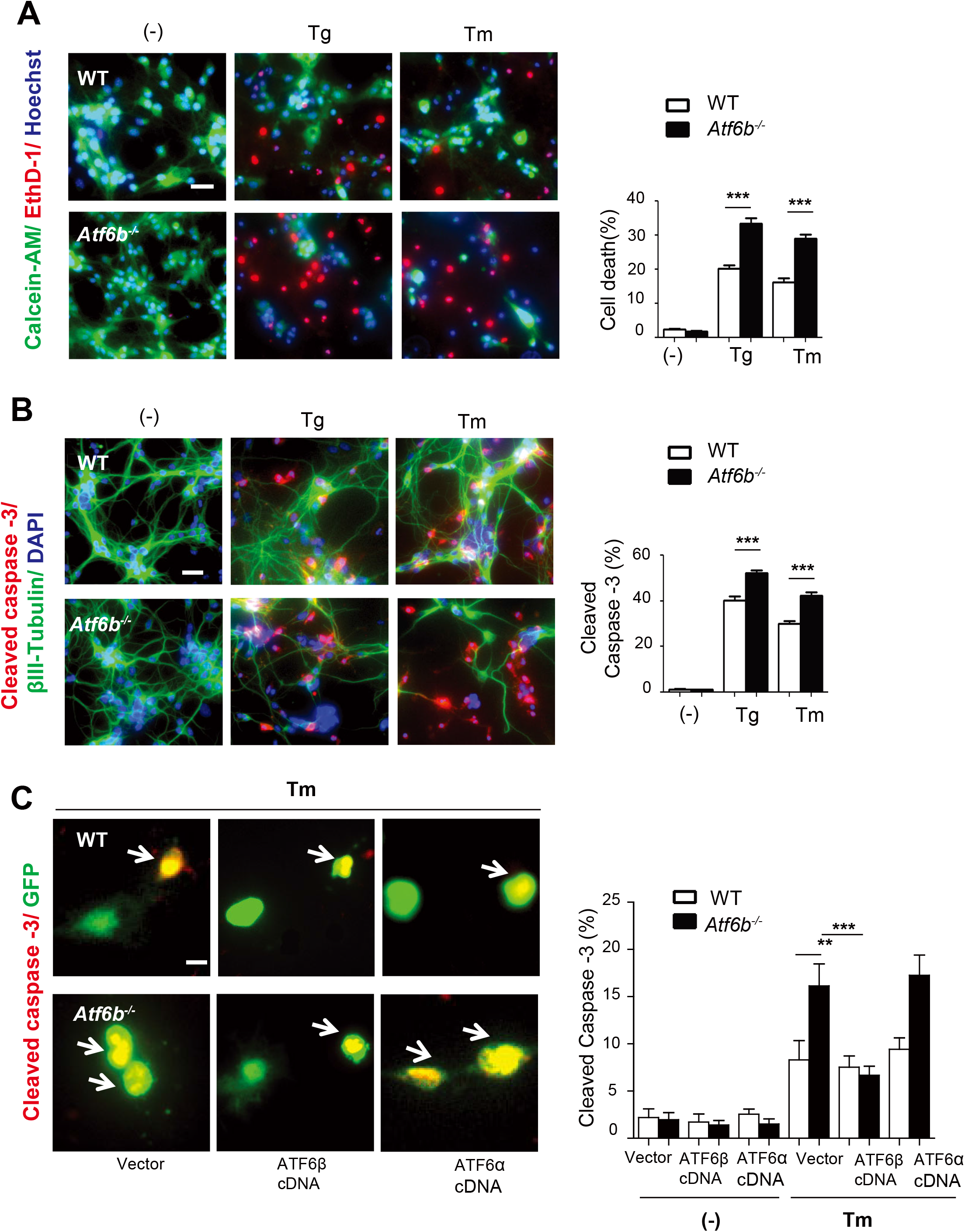
Protection of the primary hippocampal neurons by ATF6β. A, Primary hippocampal neurons were treated with Tg (300nM) or Tm (1μg/ml) for 24 h, and cell survival/death was evaluated by the LIVE/DEAD viability assay. Representative fluorescent microscopic images from four independent experiments are shown. The graph depicts the percentages of dead cells. n=4 experiments. Data are shown as mean ±SEM. ***p<0.001 by a two-way ANOVA followed by the Bonferroni test. Scale bar: 20 μm. B, Apoptosis was evaluated by immunocytochemical staining of cleaved caspase-3 (red), βIII tubulin (green) and DAPI staining (blue). Representative fluorescent microscopic images from four independent experiments are shown. The graph depicts the percentages of cleaved caspase-3-positive cells. n=4 experiments. Data are shown as mean ±SEM. ***p<0.001 by a two-way ANOVA followed by the Bonferroni test. Scale bar: 20 μm. C, Apoptosis was evaluated by immunocytochemical staining of cleaved caspase-3 (red) and GFP (green) using primary hippocampal neurons co-transfected with vector, ATF6β cDNA or ATF6α cDNA together with GFP cDNA, followed by Tm treatment for 24h. Representative fluorescent microscopic images of vector-transfected cells are shown. Arrows indicate caspase-3-positive and GFP-positive cells. The graph depicts the percentages of cleaved caspase-3-positive cells out of GFP-positive cells. n=400 cells per condition from two independent experiments. Data are shown as mean ±SEM. **p<0.01, ***p<0.001 by a two-way ANOVA followed by the Bonferroni test. Scale bar: 10 μm. Note that ATF6β cDNA, but not ATF6α cDNA, rescued *Atf6b^−/−^* neurons.

To confirm the neuroprotective role of the ATF6β-CRT axis against ER stress in hippocampal neurons, rescue experiments were performed by transfecting ATF6β and ATF6α cDNAs (Figure 5C), or by using a lentivirus-mediated CRT overexpression system (LV-CRT) (Figure 6A, B). When cells were co-transfected with ATF6β and GFP cDNAs, the number of cleaved caspase-3-positive cells in GFP-positive cells were reduced in *Atf6b^−/−^* neurons. However, this rescuing effect was not observed with ATF6α cDNA (Figure 5 C). Consistent with the rescuing effect of ATF6β, overexpression of CRT in *Atf6b^−/−^* neurons restored its expression to a similar level as that in control WT neurons (Figure 6A), and restored the survival of WT and *Atf6b^−/−^* neurons upon Tm treatment, but this effect was greater in *Atf6b^−/−^* neurons than in WT neurons (Figure 6B).

**Figure 6.**
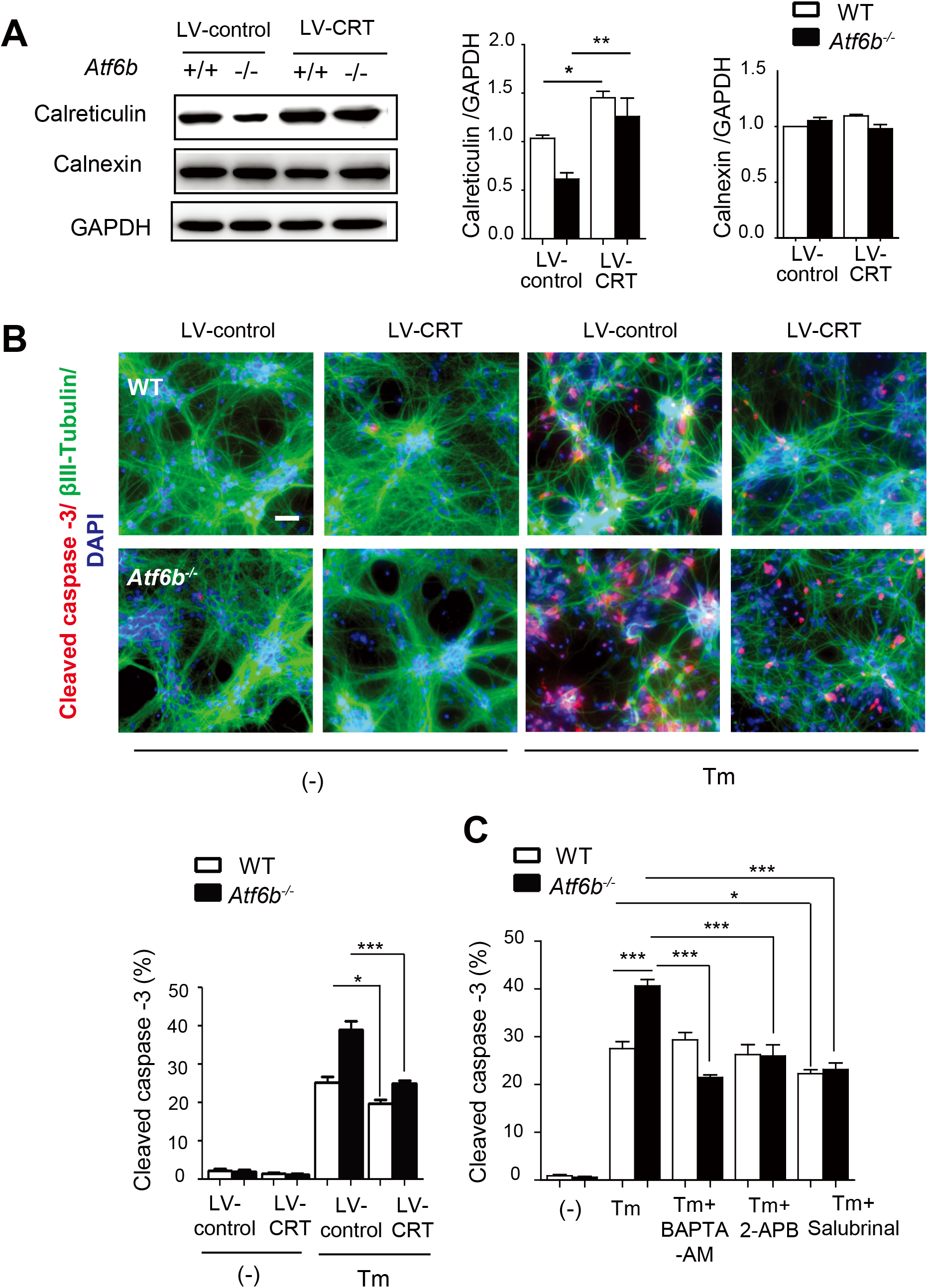
Protection of the primary hippocampal neurons by CRT overexpression or by treatment with Ca^2+^/ER stress-modulating compounds. A, B, WT and *Atf6b^−/−^* hippocampal neurons were infected with a control or CRT-expressing lentiviral vector, and the expression levels of CRT and calnexin were measured by western blotting (A). n=3 experiments. Data are shown as mean ± SEM. *p < 0.05, **p < 0.01 by a two-way ANOVA followed by the Bonferroni tests. Cells were then treated with Tm (1μg/ml) for 24 h, and cell death was evaluated by immunocytochemical staining for cleaved caspase-3 (B). Scale bar: 20 μm. n=3 experiments. Data are shown as mean ± SEM. *p < 0.05, ***p < 0.001 by a two-way ANOVA followed by the Bonferroni tests. C, WT and *Atf6b^−/−^* hippocampal neurons were treated with Tm (1μg/ml) together with BAPTA-AM (5μM), 2-APB (2μM) or salubrinal (5μM). Cell death was evaluated by immunocytochemical staining for cleaved caspase-3. n=3 experiments. Typical images are shown in Figure S5 Data are shown as mean ± SEM. *p < 0.05, **p < 0.01, ***p< 0.001 by a two-way ANOVA followed by the Bonferroni tests.

### Effects of Ca^2+^-modulating reagents and an ER stress inhibitor on ER stress-induced neuronal death

The CRT level and Ca^2+^ signaling are critical for modulating neuronal death in a neurodegeneration model(Taguchi *et al.*, 2000; Bernard-Marissal *et al.*, 2012); therefore, the effects of the Ca^2+^-modulating reagents O,O’-bis(2-aminophenyl)ethyleneglycol-N,N,N’,N’-tetraacetic acid, tetraacetoxymethyl ester (BAPTA-AM), a cell-permeable Ca^2+^ chelator, and 2-aminophosphate borate (2-APB), an inhibitor of IP3Rs and store-operated channels, and the ER stress inhibitor salubrinal were analyzed in our model. Immunocytochemical analysis revealed that both BAPTA-AM and 2-APB significantly improved survival of *Atf6b^−/−^* neurons, but not of WT neurons, under ER stress (images are shown in Figure S6 A and quantified data are shown in Figure 6C). By contrast, salubrinal enhanced survival of both WT and *Atf6b^−/−^* neurons under ER stress (images are shown in Figure S6 A and quantified data are shown in Figure 6C). These results suggest that temporal impairment of Ca^2+^ homeostasis in *Atf6b^−/−^* neurons enhances ER stress, leading to increased ER stress-induced neuronal death. Because it was reported that BAPTA-AM alone caused ER stress and neuronal death at the concentrations higher than 13μM (Paschen *et al.*, 2003), we confirmed no toxicity of BAPTA-AM and 2-APB at the concentrations in this study (5μM and 2μM, respectively) (Figure S6 B). Similarly, we analyzed the effect of salubrinal with different concentrations. Significant neuroprotection was observed at 5 and 10μM, and higher concentration (50μM or higher) enhanced ER stress-induced neuronal death in our model (Figure S6 C).

### Effect of ATF6β deletion on molecular chaperones in the ER in kainite (KA)-injected mice

Kainate (KA), an agonist of glutamate receptors, causes Ca^2+^-dependent hyperactivation of neurons, followed by the induction of ER stress and neuronal death in the hippocampus (Kezuka *et al.*, 2016). We and other groups have demonstrated the protective role of UPR signaling in KA-injected mice (Kitao *et al.*, 2001; Sokka *et al.*, 2007; Kezuka *et al.*, 2016). In this study, we investigated the role of ATF6β-CRT axis and expression of its components in these mice. RT-qPCR revealed that expression of *Atf6b*, *Calr, Canx,* and *Hspa5* mRNAs mildly, but significantly, increased after injection of KA into the mouse hippocampus (Figure 7A). Consistent with the results in cultured neurons, the level of *Calr* mRNA, but not of other mRNAs, was reduced to ~50% in *Atf6b^−/−^* mice under both sham and KA-injected conditions (Figure 7A). Western blot analysis confirmed that the level of CRT protein was decreased in the *Atf6b^−/−^* hippocampus under both sham and KA-injected conditions (Figure 7B).

**Figure 7.**
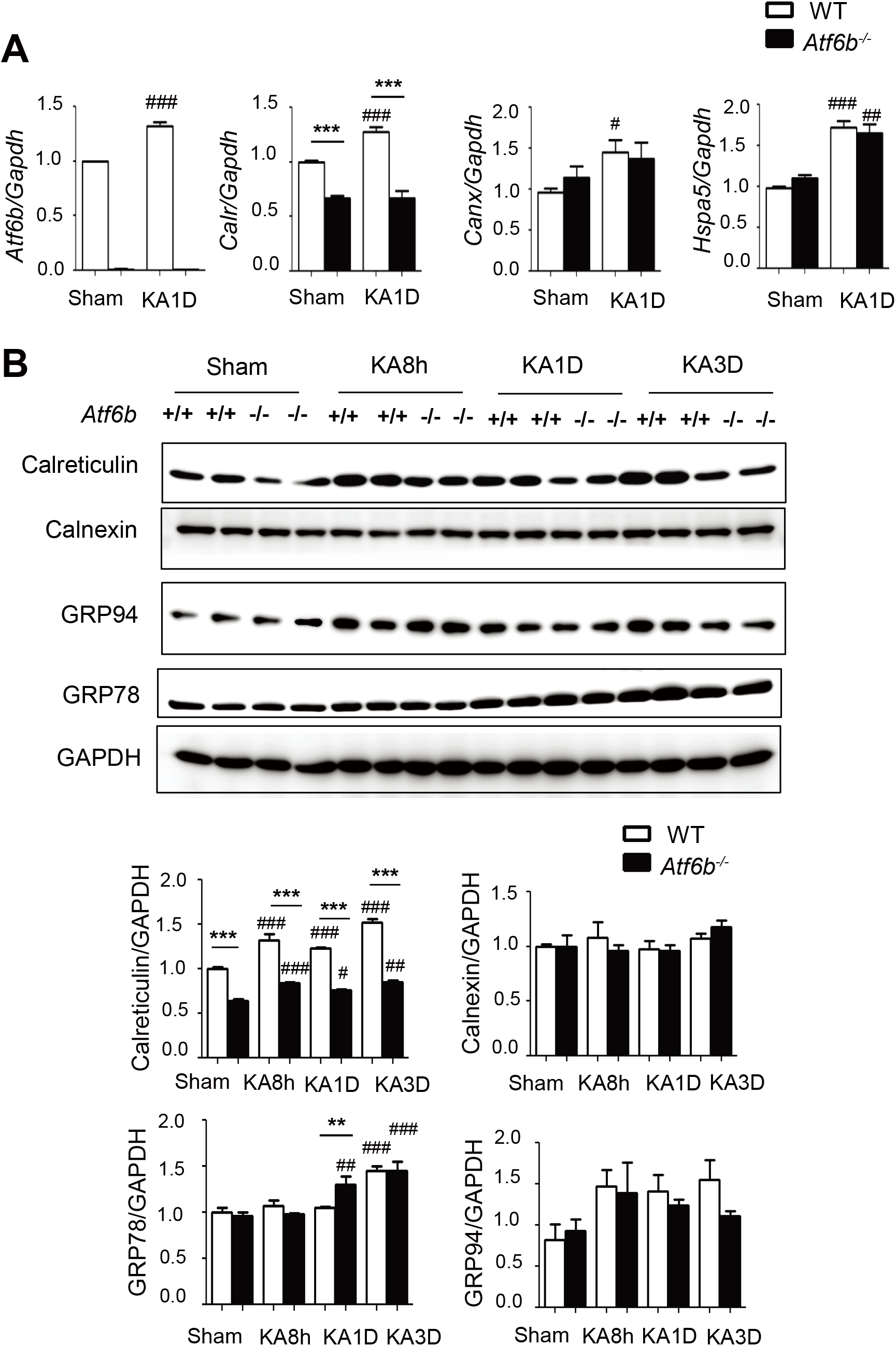
Expression of molecular chaperones in the ER in KA-injected mice. A, Total RNA was isolated from the CA3 region of hippocampi from KA-injected WT and *Atf6b^−/−^* mice and qRT-PCR was performed. n=4 mice. Data are shown as mean ± SEM. ***p < 0.001 between two genotypes and # p < 0.05, ## p < 0.01, ### p<0.001 compared to normal conditions by a two-way ANOVA followed by the Bonferroni tests. B, Protein samples were extracted from the CA3 region of hippocampi from KA-injected WT and *Atf6b^−/−^*mice and subjected to western blotting using antibodies against CRT, calnexin, GRP94 and GRP78. n=4–5 mice. Data are shown as mean ± SEM. **p < 0.001, ***p < 0.001 between two genotypes and # p < 0.05, ## p < 0.01, ### p<0.001 compared to normal conditions by a two-way ANOVA followed by the Bonferroni tests.

### The neuroprotective role of ATF6β-CRT axis against KA-induced neuronal death

The neuroprotective effect of ATF6β *in vivo* was evaluated using KA-injected mice. Consistent with our previous reports (Kitao *et al.*, 2001; Kezuka *et al.*, 2016), Nissl staining and immunohistochemical staining for cleaved caspase-3 revealed that KA caused neuronal death in the CA3 region of the hippocampus, which is one of the most KA-sensitive areas. The level of neuronal death was significantly higher in *Atf6b^−/−^* mice than in WT mice at 1 and 3 days after KA injection (Figure 8A, B). To analyze the involvement of CRT in KA-induced neuronal death, *Calr^+/−^* mice, which developed normally and showed no gross phenotypes with a reduced level of *Calr* expression (Figure S7), were injected with KA and neuronal death was evaluated. Consistent with the results obtained with *Atf6b^−/−^* mice, the level of neuronal death in the hippocampus was significantly higher in *Calr^+/−^* mice than in WT mice (Figure 8C).

**Figure 8.**
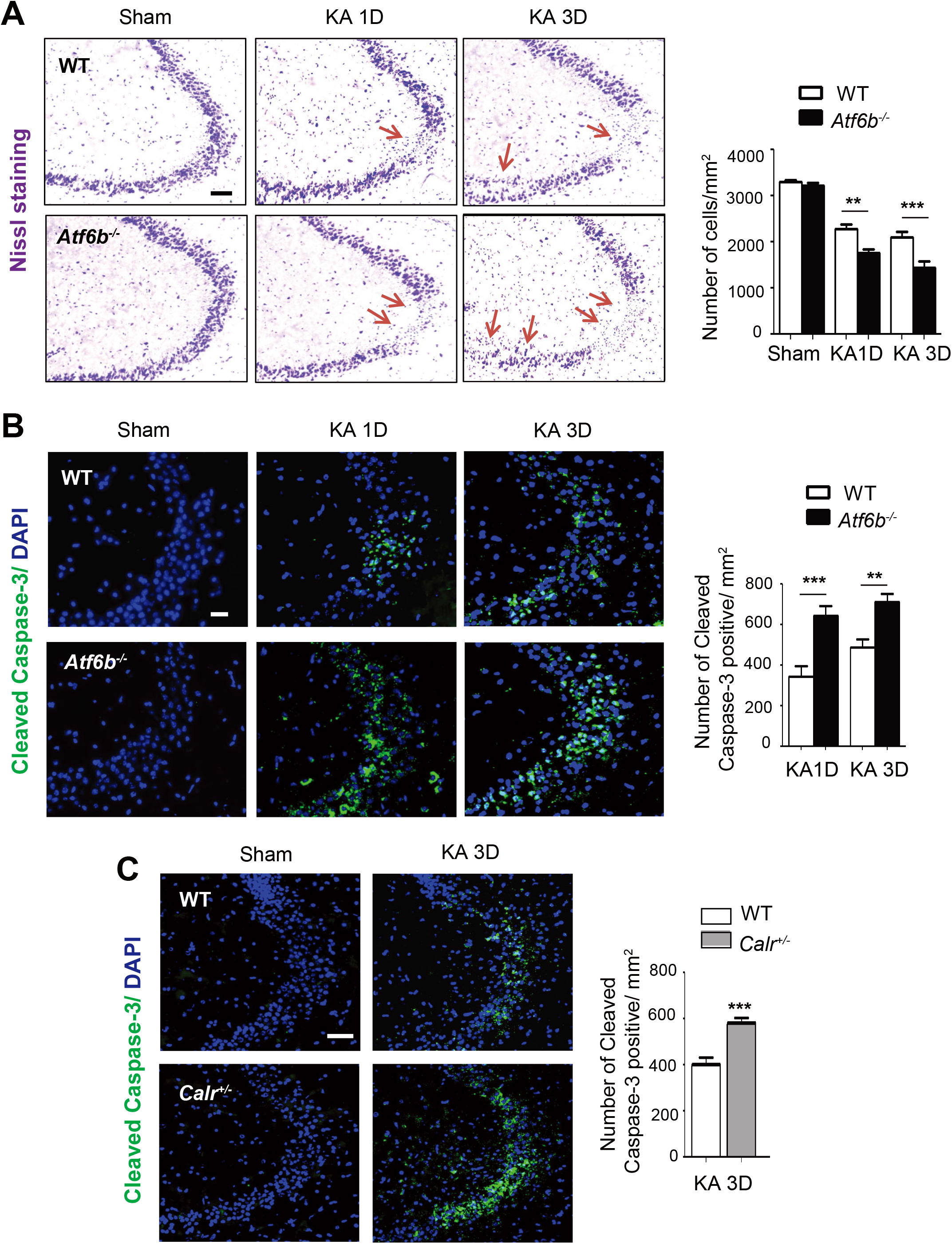
Neuroprotection by ATF6β and CRT in KA-injected mice. A, Brain sections containing the CA3 area of the hippocampus from WT and *Atf6b^−/−^* mice were subjected to Nissl staining. The right graph depicts the number of surviving CA3 neurons. n=6-8 mice. Data are shown as mean ± SEM. **p < 0.01, ***p < 0.001 by a two-way ANOVA followed by the Bonferroni tests. Scale bar: 100 μm. B, C, Brain sections including the CA3 area of the hippocampus from WT and *Atf6b^−/−^* mice (B) (n=6–7 mice) or WT and *Calr^+/−^* mice (C) (n=6 mice) were subjected to immunohistochemical staining for cleaved caspase-3. The right graphs depict the number of cleaved caspase-3-positive CA3 neurons. Data are shown as mean ± SEM. **p < 0.01, ***p < 0.001 by a two-way ANOVA followed by the Bonferroni tests in (B) and by the Mann-Whitney U test in (C). Scale bar: 50 μm.

### ATF6β-mediated regulation of neuronal activity and Ca^2+^ homeostasis after KA administration

To elucidate the mechanism underlying the enhanced level of neuronal death in *Atf6b^−/−^* hippocampus after KA injection, earlier events following KA injection were investigated. qRT-PCR (Figure S8 A) and immunohistochemistry (Figure S8 B) revealed that expression of the immediate-early genes such as *Fos* (c-Fos), *Fosb* and *Bdnf* was induced in both genotypes after KA-injection, but was higher in *Atf6b^−/−^* mice, suggesting that hyperactivity is involved in the enhanced level of neuronal death in the *Atf6b^−/−^* hippocampus. The effects of the Ca^2+^-modulating reagent 2-APB and the ER stress inhibitor salubrinal were next analyzed. They did not cause neuronal death at the doses used in this study (Figure S8 C). Immunohistochemical analysis revealed that both reagents significantly improved neuronal survival in the *Atf6b^−/−^* hippocampus after KA injection (Figure 9A, B), suggesting that temporal dysregulation of Ca^2+^ homeostasis in *Atf6b^−/−^* neurons enhances ER stress, leading to increased ER stress-induced neuronal death.

**Figure 9.**
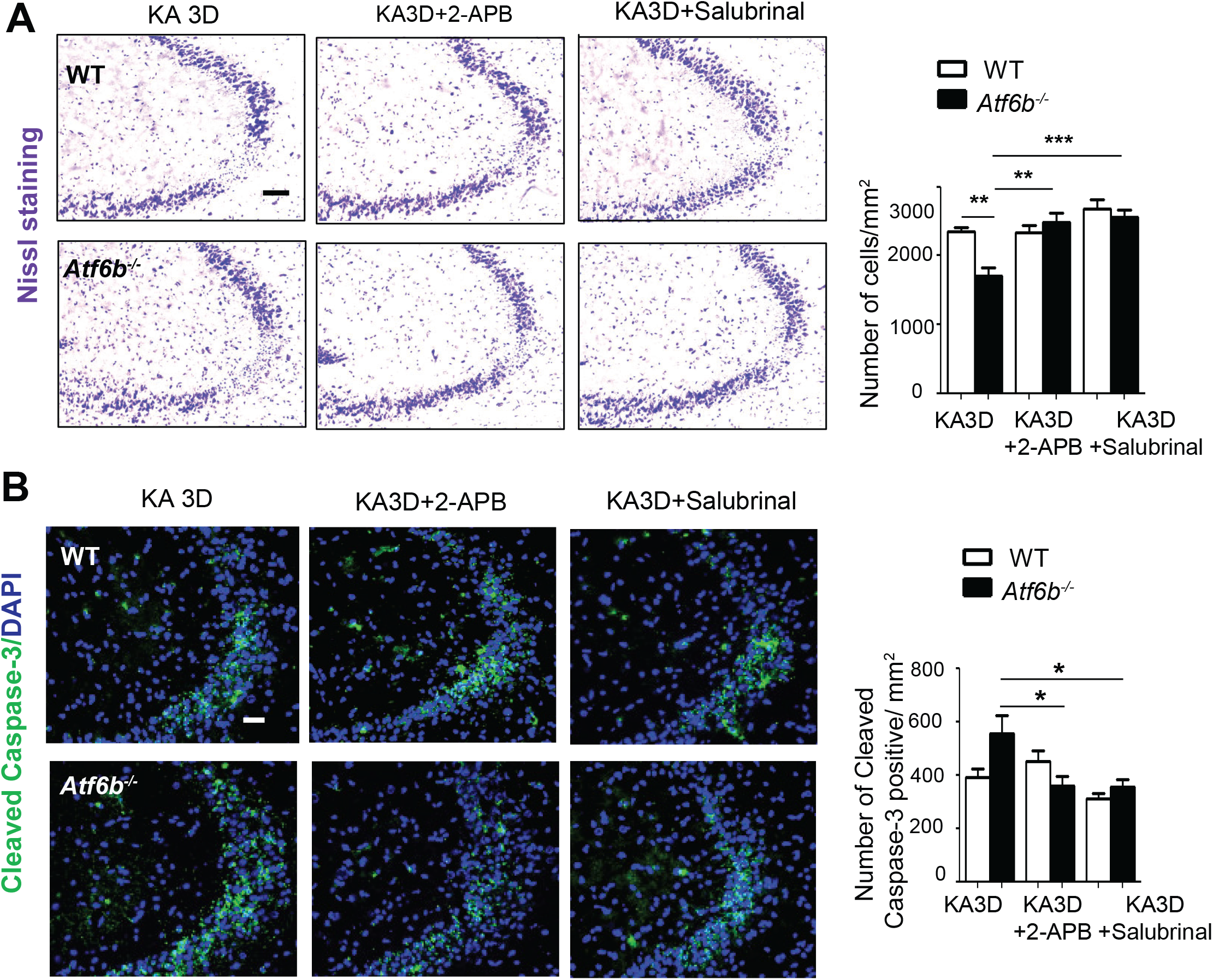
Ca^2+^ dysregulation in *Atf6b^−/−^* mice after KA-injection. Brain sections including the CA3 area of the hippocampus obtained from WT and *Atf6b^−/−^* mice at 3 days after injection with KA, KA plus 2-APB and KA plus salubrinal were subjected to Nissl staining (A) or immunohistochemical staining for cleaved caspase-3 (B). The right graphs depict the number of surviving CA3 neurons (A) and cleaved caspase-3-positive cells (B), respectively. n=6 mice. Data are shown as mean ± SEM. *p < 0.05, **p < 0.01, ***p < 0.001 by a two-way ANOVA followed by the Bonferroni tests.

## Discussion

The major findings of the current study are that ATF6β specifically regulates CRT expression in the CNS and that the ATF6β-CRT axis plays an important role in survival of hippocampal neurons upon exposure to ER stress and excitotoxicity.

CRT is a Ca^2+^-binding molecular chaperone in the ER that functions in diverse cellular processes such as Ca^2+^ homeostasis, protein folding, gene expression, adhesion, and cell death (Michalak *et al.*, 2009; Wang *et al.*, 2012). It is also important for organogenesis especially in the heart, brain, and ventral body wall (Rauch *et al.*, 2000). Deficiency of CRT leads to defects in myofibrillogenesis and thinner ventricular walls in the heart (Mesaeli *et al.*, 1999). Interestingly, overexpression of CRT in the heart also causes severe phenotypes such as arrhythmias and sudden heart block following birth (Nakamura *et al.*, 2001). Therefore, the transcription of CRT in the heart is strictly controlled by several transcriptional factors such as Nkx2.5, COUP-TF1, GATA6 and Evi-1 (Qiu *et al.*, 2008). CRT is also highly expressed in the developing brain and retina, and its deficiency leads to a defect in closure of neural tubes (Rauch *et al.*, 2000). Although it is unclear whether overexpression of CRT is toxic in the CNS, it is possible that the similar strict regulation of CRT expression is required and that ATF6β is utilized in addition to ATF6α for this purpose.

Our results indicate an important, but a bit puzzling role for ATF6β in CRT expression in the CNS. All the data from RNA-sequence to the promoter analysis suggested that CRT expression was ATF6β-dependent in primary hippocampal neurons. However, overexpression of ATF6α and ATF6β both enhanced CRT promoter activity, and those effects were severely diminished when the CRT promoter contained mutated ERSEs (Figure 3B, Figure S3). These results were consistent with a previous report demonstrating that both homo- and heterodimer of ATF6α and ATF6β bind to ERSEs in a similar manner (Yoshida *et al.*, 2001). In fact, the amino acid sequences of the basic regions of ATF6α and ATF6β, which are required for DNA binding, are 91% identical (21 of 23 amino acid) (Yoshida *et al.*, 1998). In contrast to CRT expression, those of other molecular chaperones in the ER such as GRP78 and GRP94 were more ATF6α-dependent (Figure 2A) and the deletion of ATF6β only temporally affected their expressions under ER stress (Figure 4A). These results may raise a scenario that, in the CNS, expression of molecular chaperones in the ER is generally governed by ATF6α as previously described (Yamamoto *et al.*, 2007) and that ATF6β functions as a booster if their levels are too low. However, expression of CRT is somewhat governed by ATF6β, and ATF6α functions as a booster. The underlying mechanism for this scenario is not clear yet, but neurons may require a high level of CRT expression even under normal condition, as described in Table S2, which may lead to the development of a unique biological system to constitutively produce CRT in neurons. Further studies are required to clarify the molecular basis how this unique system is constructed and regulated.

The current study demonstrated that ATF6β has neuroprotective roles against ER stress and KA-induced neurotoxicity, both of which are associated with CRT. Deletion of *Atf6b* reduced CRT expression (Figure 4A, B), impaired intracellular Ca^2+^ homeostasis (Figure 4C), and enhanced ER stress-induced death of cultured hippocampal neurons (Figure 5) and Neuro 2a cells (Figure S5). Overexpression of ATF6β or CRT, but not ATF6α, rescued *Atf6b^−/−^* hippocampal neurons against ER stress-induced cell death (Figure 5C, Figure 6A, B). The lack of rescuing effect of ATF6α may be due to the fact that this molecule enhances the expression of different genes including cell death-related molecule CHOP in addition to molecular chaperons in the ER (Yoshida *et al.*, 2000). Consistent with the role of ATF6β-CRT axis in the Ca^2+^ homeostasis and neuroprotection, treatment with the Ca^2+^-modulating reagents BAPTA-AM and 2-APB or the ER stress inhibitor salubrinal restored the survival of neuronal cells under ER stress conditions (Figure 6C, Figure S6 A, C). Interestingly, the effects of the Ca^2+^-modulating reagents were observed only in *Atf6b^−/−^* neurons, while that of the ER stress inhibitor was observed in both WT and *Atf6b^−/−^* neurons (Figure 6C). These results suggest that the ATF6β-CRT axis maintains intracellular Ca^2+^ levbel, which contributes to suppression of ER stress and prevention of apoptosis. Similarly, our *in vivo* results suggest that a reduced level of CRT, Ca^2+^-mediated neuronal hyperactivity, and subsequent ER stress underlie enhanced neuronal death in the hippocampus of KA-injected *Atf6b^−/−^* mice (Figure 8 and Figure 9).

Accumulating evidence suggests that a reduced level of CRT is associated with the pathologies of neurodegenerative diseases such as amyotrophic lateral sclerosis (ALS) (Bernard-Marissal *et al.*, 2012; Bernard-Marissal *et al.*, 2015) and Alzheimer’s disease (AD) (Taguchi *et al.*, 2000; Lin *et al.*, 2014). In a mutant superoxide dismutase (mSOD1) model of ALS, activation of the Fas/nitric oxide (NO) pathway reduces CRT expression in motoneurons, which further activates Fas/NO signaling on one hand and enhances ER stress and neuronal death on the other hand (Bernard-Marissal *et al.*, 2012). Consequently, the level of CRT is drastically decreased in 50% of fast fatigable motoneurons (Bernard-Marissal *et al.*, 2015). Although it is unclear whether ATF6β is involved in the Fas/NO-mediated reduction of the CRT expression, it will be intriguing to analyze the expression profile of ATF6β and the phenotype of *Atf6a^−/−^* mice in these models.

Consistent with the observations in ALS, the level of CRT is reduced in the brains (Taguchi *et al.*, 2000) and sera (Lin *et al.*, 2014) of AD patients, raising the possibility that CRT is a good biomarker of AD (Lin *et al.*, 2014). However, it was also reported that a portion of CRT is located on the cell surface membrane, and acts as a receptor for C1q, the recognition subunit of the first component of complement. The C1q-CRT complex induces oxidative neurotoxicity (Luo *et al.*, 2003); therefore, CRT may also have a pathological role in AD. Further analysis is required to elucidate the precise role of CRT and involvement of ATF6β in AD.

Although the function of ATF6β has been considered to be very limited or redundant compared with that of ATF6α, our results emphasize the critical and beneficial roles of ATF6β in neuropathological conditions. A recent study also demonstrated that ATF6β is functional in the heart, especially during the pressure overload-induced cardiac hypertrophic response (Correll *et al.*, 2019). The role of ATF6β may be determined by the need for specific molecular chaperones in the ER such as CRT, which may differ between tissues. Further studies dissecting the cell- and tissue-specific roles of ATF6β will help to elucidate the function of the UPR in pathophysiological conditions.

## Materials and Methods

### Animals

All animal experiments were conducted according to the guidelines of the Animal Care and Use Committee of Kanazawa University (Approval No. AP-184013). *Atf6a^+/−^* and *Atf6b^+/−^* mice were generated as previously described (Yamamoto *et al.*, 2007), and backcrossed with the C57BL/6 strain for more than eight times at the Institute of Laboratory Animals, Graduate School of Medicine, Kyoto University. *Calr^+/−^* mice were generated as previously described (Tokuhiro *et al.*, 2015), and provided by the RIKEN BioResource Research Center (Tsukuba, Ibaraki, Japan). *Atf6a^+/−^* and *Atf6b^+/−^* mice were intercrossed to obtain wild-type (WT), *Atf6a^−/−^*, and *Atf6b^−/−^* mice. These lines were maintained by mating mice of the same genotype at the Institute for Experimental Animals, Advanced Science Research Center, Kanazawa University. *Calr^+/−^* mice were maintained by mating mice with WT mice in the C57BL/6 background. WT, *Atf6a^−/−^*, *Atf6b^−/−^*, and *Calr^+/−^* mice (male; age, 10– 12 weeks; weight, 25–30 g) were used for experiments.

To develop a KA injection model, mice were anesthetized, and saline or KA (0.2 μg/μl, 0.5 μl in total; Sigma, St Louis, MO, USA) was injected unilaterally into the hippocampus (from Bregma: dorso-ventral,-2.0; medio-lateral, −2.4; anterior-posterior, −1.8), as previously described (Kezuka *et al.*, 2016). In some cases, 2-APB (12 μM, 0.5 μl in total; FUJFILM Wako Pure Chemical Co., Osaka, Osaka, Japan) or salubrinal (1mg/kg; Cayman Chemical, Ann Arbor, MI, USA) was co-injected with KA into the hippocampus, or intraperitoneally injected 30 min before KA administration, as previously described (Sokka *et al.*, 2007; Kim *et al.*, 2014; Ikebara *et al.*, 2017). Mice were sacrificed at the indicated timepoints after KA injection, and brain samples were prepared for histological and biochemical analysis.

### Cell cultures

Primary hippocampal and cortical neurons were isolated from embryonic day 17.5 (E17.5) WT and *Atf6b^−/−^* mice, as previously described, with minor modifications (Kaech & Banker, 2006). Briefly, hippocampi and cerebral cortices were harvested from prenatal mice, and digested using neuron dissociation solution (FUJFILM Wako Pure Chemical Co.). After isolation, neurons were plated into 24-well culture plates precoated with poly-L-lysine (10μg/ml; Sigma) at a density of 8×10^5^ cell/well, and cultured in Neurobasal Medium (Life Technologies, Carlsbad, CA, USA) supplemented with 2% B-27 serum free supplement (Life Technologies), 0.4 mM L-glutamine (Sigma), 5% fetal bovine serum (FBS)(Sigma), 100U/ml penicillin and 100 μg/ml streptomycin (Nacalai Tesque, Kyoto, Kyoto, Japan). After 3 days, neurons were used for experiments. Hippocampal neurons were treated with the ER stressors Tg (300nM; Sigma), DTT (1mM; Nacalai Tesque) and Tm (1μg/ml; FUJFILM Wako Pure Chemical Co.). In some cases, they were treated with BAPTA-AM (5μM; Dojindo Molecular and Technologies Inc., Mashiki-machi, Kumamoto, Japan), 2-APB (2μM) or salubrinal (5μM) in addition to Tm for the indicated durations.

Astrocytes were isolated from the cerebral cortex of postnatal day 1–3 WT mice, as previously described (McCarthy & de Vellis, 1980), and cultured in Dulbecco’s Modified Eagle Medium (DMEM) supplemented with 10% FBS and penicillin/streptomycin. Cells were used for experiments after achieving full confluency.

MEFs were isolated from the skin of E15.5 WT and *Atf6b^−/−^* mice, as previously described (Yamamoto *et al.*, 2007), and were cultured in DMEM supplemented with 20% FBS and penicillin/streptomycin. Cells were used for experiments after achieving full confluency.

Neuro 2a cells were plated at a density of 5 ×10^4^ cells/well in 24- or 12-well culture plates, and cultured in DMEM supplemented with 10% FBS and penicillin/streptomycin. Cells were used for experiments after achieving 70% of confluency.

### Preparation and transfection of plasmids

Plasmids expressing full length mouse ATF6α and ATF6β were constructed by inserting ATF6α and ATF6β cDNAs into the pCDF1-MCS2-EF1-Puro expression vector (System Biosciences, Palo Alto, CA, USA). pcDNA3.1(+) GFP was obtained from Invitrogen/Thermo Fisher Scientific (Waltham, MA, USA). CAT plasmids containing the mouse CRT promoter (pCC1 and pCC3) were provided by Dr. Marek Michalak (University of Alberta) (Waser *et al.*, 1997). Both plasmids contain ERSE, a consensus of CCAATN_9_CCACG (Yoshida *et al.*, 1998) (Figure 3A). Luciferase plasmids containing huCRT(wt) and huCRT(mut), with the two ERSEs mutated in the latter, and a plasmid containing the WT human GRP78 promoter (huGRP78) were constructed as previously described (Yoshida *et al.*, 1998). The pRL-SV40 plasmid was obtained from Promega (Madison, WI, USA). Plasmids for Ca^2+^ imaging such as pCMV G-CEPIA1er, pGP-CMV-GCaMP6f and pCMV CEPIA2mt were obtained from Addgene (Watertown, MA, USA). Cells were transfected with each plasmid for 5 h using Lipofectamine 2000 (Life Technologies) and further incubated for 24-48 h. In our model, transfection efficiency was approximately 5% in primary neurons.

### Preparation and infection of lentivirus vectors

The lentivirus vector expressing full-length mouse CRT under the control of the human eukaryotic translation elongation factor 1 α1 promoter and the lentivirus vector alone was purchased from VectorBuilder (Chicago, IL, USA). Viral stocks had titers of ~10^9^ plaque-forming units/ml. Hippocampal neurons were infected with the CRT-expressing (LV-CRT) or control (LV-control) lentivirus vector at a multiplicity of infection 10 for 16 h and further incubated for 48-72 h.

### Preparation and transfection of ATF6β-targeting siRNAs

ATF6β-specific siRNAs, namely, ATF6β-siRNA1 (SASI_Mm01_00110468) and ATF6β-siRNA2 (SASI_Mm01_00110470), and control-siRNA were obtained from Sigma. Neuro2a cells were transfected with each siRNA for 5 h using Lipofectamine RNAiMAX (Life Technologies) and further incubated for 24-48 h.

### qRT-PCR

Total RNA was extracted from the indicated mouse tissues and cultured cells using RNeasy^®^ Lipid Tissue Mini Kit (Qiagen, Valencia, CA, USA). Reverse transcription reactions containing 1 μg of total RNA were performed using PrimeScript™ (Takara, Otsu, Shiga, Japan). Individual cDNAs were amplified with THUNDERBIRD™ SYBR qPCR Mix (TOYOBO CO, LTD, Osaka, Osaka, Japan) using specific primers for *Atf6b*, *Atf6a*, *Calr*, *Canx*, *Hspa5*, *Hsp90b1*, *Fos*, *Fosb*, *Bdnf* and *Gapdh*. The primers are listed in Table S3. The comparative Ct method was used for data analyses with MxPro 4.10 (Agilent Technologies, Santa Clara, CA, USA). Values for each gene were normalized against the *Gapdh* expression level.

### Western blotting

Samples from the indicated mouse tissues and cultured cells were solubilized in RIPA buffer, which contained 10 mM Tris (pH 7.6), 1 mM EDTA, 150 mM NaCl, 1% NP-40, 0.1% SDS, 0.2% sodium deoxycholate, 1 mM PMSF, 1μg/ml aprotinin, 10 mM NaF, and 1 mM Na_3_VO_4_. To detect endogenous ATF6β protein in primary hippocampal neurons, RIPA buffer without sodium deoxycholate was used. Membranes were incubated with 3% bovine serum albumin for 1 h and then with the primary antibodies for overnight at 4℃. The primary antibodies included those against ATF6β (853202; Biolegend, San Diego, CA, USA; 1:500), CRT (SPA-600; Enzo Life Sciences Inc., Farmingdale, NY, USA; 1:2000 and 10292-1-AP; Proteintech, Rosemont, IL, USA; 1:1000), calnexin (SPA-865; Enzo Life Sciences Inc.; 1:1000), KDEL (PM-059; Medical & Biological Laboratories, Nagoya, Aichi, Japan; 1:1000), and GAPDH (016-25523; FUJFILM Wako Pure Chemical Co.; 1:2000). Sites of primary antibody binding were determined using an enhanced chemiluminescence system (GE Healthcare, Pittsburgh, PA, USA). Horseradish peroxidase (HRP)-conjugated secondary antibodies (Santa Cruz Biotechnology, Dallas, TX, USA) were used to detect immunoglobulins from mouse or rabbit. The intensity of each band was, quantified by using Image J software (version 1.50i, Wayne Rasband, National Institutes of Health, Bethesda, MD, USA, https://imagej.nih.gov/ij/).

### Histology and immunohistochemistry

At the indicated timepoints after KA injection, mice (WT, Atf6b^−/−^, and Calr^+/−^) were deeply anesthetized with isoflurane and transcardially perfused with phosphate-buffered saline (PBS) followed by 4% paraformaldehyde prepared in 0.1 M phosphate buffer (pH 7.4). Brains were harvested, post-fixed with 4% paraformaldehyde for 8 h, and cryoprotected in 30% sucrose for at least 24 h. Cortical sections (10 μm-thick coronal sections containing the hippocampus (between Bregma −1.5 and −2.1 mm)) were cut on a cryostat (Leica Biosystems, Wetzler, Germany). Sections were processed for Nissl staining (Cresyl violet staining) or immunohistochemistry with antibodies against cleaved caspase-3 (Asp175; Cell Signaling Technology, Inc. Danvers, MA, USA; 1:500) and c-Fos (PC05; Merck; 1:200). Nuclei were visualized with 4’,6-diamidino-2-phenylindole (DAPI; Sigma). Anti-rabbit and anti-mouse Alexa Fluor 488-conjugated (Life Technologies; 1:200) and Cy3-conjugated (Jackson ImmunoResearch Laboratories, Inc., West Grove, PA, USA; 1:200) secondary antibodies were used to visualize immunolabeling. Imaging was performed using a laser scanning confocal microscope (Eclipse TE200U; Nikon, Tokyo, Japan) and Nikon EZ-C1 software or using a light and fluorescence microscope (BZ-X700; KEYENCE, Osaka, Japan).

### In situ hybridization-immunohistochemistry

*In situ* hybridization was performed as previously described (Hattori *et al.*, 2014). In brief, a ~600 bp mouse ATF6β cDNA fragment was PCR-amplified using 5’-AACAGGAAGGTTGTCTGCATCAT-3’ and 5’-GTATCCTCCCTCCGGTCAAT-3’ primers and inserted into the pGEM-T vector (Promega). The plasmid was linearized using EcoRV and ApaI to synthesize the antisense and sense probe, respectively. Brains were removed from mice after perfusion with PBS and immediately placed at −80℃. Serial 14 μm-thick coronal sections were obtained using a cryostat and hybridized with a digoxigenin-labeled ATF6β RNA probe. After development and thorough washing with PBS, brain sections were subjected to immunohistochemistry using a mouse anti-NeuN antibody (MAB377; Merck, Kenilworth, NJ, USA; 1:500) followed by incubation with an anti-mouse IgG antibody (Vector Laboratories, Inc., Burlingame, CA, USA). The sections were developed in peroxidase substrate solution (ImmPACT DAB, Vector Laboratories, Inc.). Imaging was performed using a light and fluorescence microscope (BZ-X700, KEYENCE).

### RNA-sequencing

Total RNA purified from hippocampi of WT and *Atf6b^−/−^* mice (n=2 per group) was used to prepare RNA libraries using a TruSeq Stranded mRNA Sample Preparation Kit (Illumina, Inc., San Diego, CA, USA), with polyA selection for ribosomal RNA depletion. The RNA libraries were generated from 500 ng of total hippocampal RNA and sequenced on an Illumina HiSeq 2000 to obtain paired-end 101 bp reads for each sample.

RNA-sequencing reads were mapped to the Mouse genome (GRCm.38.p6/mm10) using STAR v2.7.0f (Dobin *et al.*, 2013). Aligned reads were counted and assigned to genes using Ensembl release 99 gene annotation (Cunningham *et al.*, 2019). Gene expression levels were quantified using Cufflinks v2.2.1 (Trapnell *et al.*, 2013), and denoted by fragments per kilobase of exon per million reads mapped (FPKM) values, which were normalized by the number of RNA fragments mapped to the reference genome and the total length of all exons in the respective transcripts. Differentially expressed genes between WT and *Atf6b^−/−^* hippocampi were identified by Cuffdiff, which is part of the Cufflinks toolkit. Two replicates per group were combined as the input of Cuffdiff, and q- and p-values were reported to show the significance of differentially expressed genes. The raw reads are available in the DNA Data Bank of Japan (DDBJ) with DDBJ Sequence Read Archive (DRA) accession number, DRA011345.

### Ca^2+^ measurement in the intracellular organelles and in the cytosol

Ca^2+^ levels in the ER, cytosol and mitochondria of hippocampal neurons were measured using G-CEPIA1er, GCaMP6f and CEPIA2mt, respectively. Forty-eight hours after transfection, Ca^2+^ imaging was performed using a light and fluorescence microscope (BZ-X700, KEYENCE) under normal and ER stress conditions. The intensity of the fluorescence in each cell was measured using ImageJ software for 200 neurons in each condition.

### Immunocytochemistry

Cultured hippocampal neurons and Neuro 2a cells were fixed in 4% paraformaldehyde for 15 min at room temperature, and permeabilized in 0.3% Trinton-X100 for 10 min. Primary antibodies against cleaved caspase-3 (Asp175, Cell Signaling Technology, Inc; 1:800) and βIII tubulin (MAB1637; Merck; 1:500) were used. Nuclei were visualized with DAPI (Sigma). Imaging was performed using a light and fluorescence microscope (BZ-X700, KEYENCE).

### LIVE/DEAD viability assay

Living and dead neurons and Nuro 2a cells were evaluated using a LIVE/DEAD Viability Assay kit (Life Technologies). In brief, cells were washed in PBS and incubated with calcein-AM (1μM), EthD-1 (2μM) and Hoechst 33342 (1 μg/ml, Dojindo Molecular and Technologies Inc.) in regular medium. Imaging was performed using a fluorescence microscope (BZ-X700, KEYENCE).

### Reporter assay

#### Luciferase assay

At 48 h after transfection, cells were lysed in 100 μl of Passive Lysis Buffer (Promega). Firefly luciferase and Renilla luciferase activities were measured using the Dual-Luciferase Reporter Assay System (Promega) and analyzed as previously describe (Yoshida *et al.*, 1998).

#### CAT ELISA

At 48 h after transfection, cells were lysed in 125 μl of lysis buffer provided with a CAT ELISA kit (Sigma). CAT expression and Renilla luciferase activities were measured and the ratio was calculated.

### Image quantification

For the LIVE/DEAD viability assay and immunocytochemistry, four images per well were acquired and the numbers of EthD-1-positive cells/Hoechst 33342-positive cells and cleaved caspase-3-positive cells/DAPI-positive total cells were counted using ImageJ software, respectively. For Nissl staining, three brain sections containing the hippocampal CA3 area close to the KA injection site were selected per mouse and the number of Nissl-positive neurons was counted using ImageJ software. For immunohistochemistry, two brain sections with the highest numbers of cleaved caspase-3-positive cells and c-Fos-positive cells were selected per mouse, and the numbers of these cells were counted using ImageJ software.

### Statistical analyses

Statistical analyses were performed using the Mann-Whitney U test or a one-way or two-way analysis of variance (ANOVA) followed by the Tukey/Bonferroni test. GraphPad Prism software 5.0 was used for statistical analyses. A p-value less than 0.05 was considered statistically significant.

## Acknowledgements

We are grateful to Dr. Marek Michalak (University of Alberta) for providing the CAT plasmid containing the mouse CRT promoters (pCC1 and pCC3). We also thank Dr. Masahito Ikawa (Osaka University) for providing *Calr^+/−^* mice. This work was supported by a Grant-in Aid for Scientific Research (18K06500 to OH) from the Ministry of Education, Science, Technology, Sports and Culture of Japan, by a Grant (20gm1410005s0301) from the Japan Agency for Medical Research and Development (AMED), and by Kanazawa University SAKIGAKE Project 2018 and the CHOZEN project.

